# Protein folding modulates the adhesion strategy of Gram positive pathogens

**DOI:** 10.1101/743393

**Authors:** Alvaro Alonso-Caballero, Daniel J. Echelman, Rafael Tapia-Rojo, Shubhasis Haldar, Edward C. Eckels, Julio M. Fernandez

**Affiliations:** Department of Biological Sciences, Columbia University, NY 10027, USA

## Abstract

Gram positive bacteria colonize mucosal tissues against large mechanical perturbations, such as coughing, which generate large shear forces that exceed the ability of non-covalent bonds to remain attached. To overcome these challenges, the pathogen *Streptococcus pyogenes* utilizes the protein Cpa, a pilus tip-end adhesin equipped with a Cys-Gln thioester bond. The reactivity of this bond towards host surface ligands enables covalent anchoring of the bacterium, allowing it to resist large mechanical shocks; however, colonization also requires cell migration and spreading over surfaces. The molecular mechanisms underlying these seemingly incompatible requirements remain unknown. Here, we demonstrate a magnetic tweezers force spectroscopy assay that resolves the dynamics of Cpa thioester bond under force. While folded at forces < 6 pN, Cpa thioester bond reacts reversibly with amine ligands, of common occurrence in inflammation sites; however, mechanical unfolding and exposure to forces higher than 35 pN blocks thioester reactivity entirely. We propose that this folding-coupled thioester reactivity switch allows the adhesin to hop and sample host surface ligands at low force (nomadic mobility phase), and yet gets covalently anchored in place while under mechanical stress (locked phase). We dub such bonds “smart covalent bonds”, adding a novel class to the known repertoire of non-covalent adhesion strategies that include slip bonds, and catch bonds.

## Introduction

In the ancient arms race between host and pathogen, bacteria have evolved novel adhesion strategies such as biofilm formation^1,2^, non-covalent catch bond binding^3,4^, and direct covalent binding to host substrates^5,6^. In particular, Gram positive bacteria express a class of protein adhesins that contain internal Cys-Gln thioester bonds^5–7^. The thioester bond functions as an electrophilic substrate to draw a nucleophilic ligand, creating a covalent crosslink between a ligand and the adhesin of the bacterium^6^. Thioester bonds have evolved to permit bacterial adherence under large mechanical stresses^8^; however, bacterial colonization also benefits from cell rolling and spreading over surfaces^9,10^, and the molecular mechanisms reconciling the interplay between mobility and covalent anchoring are not known.

We have recently demonstrated a novel assay to study the reactivity of the pilus-tip thioester adhesin Cpa from the Gram positive pathogen *Streptococcus pyogenes*^11^ (**Figure 1a**), the causative agent of strep throat and the necrotizing fasciitis^12^. Similar to our assays for disulfide bond mechano-chemistry^13-15^, our Atomic Force Microscopy (AFM) force spectroscopy assay directly measured the presence or absence of the thioester bond in unfolding Cpa adhesins. However, our assay could not probe how force regulated Cpa folding and its coupling with thioester dynamics. AFM force spectroscopy is limited to operating at high forces and over short times^16^, so probing thioester dynamics in relation to the folding dynamics of its parent Cpa protein required a different approach.

**Figure 1.**
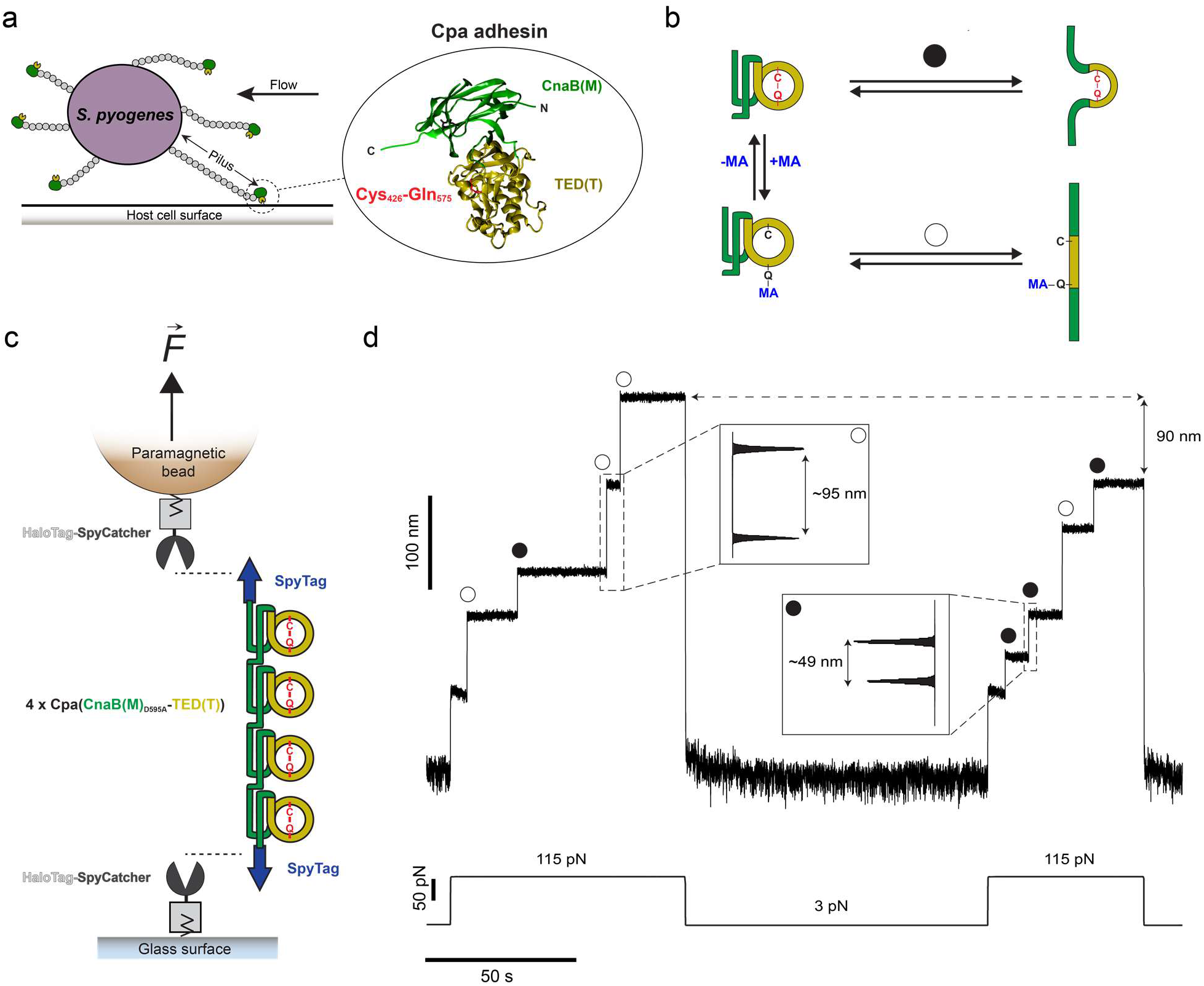
Mechano-chemistry of S. *pyogenes* Cpa adhesin. **a)** *S. pyogenes* attach to host cell surfaces through the Cpa protein, present in the tip-end of the pili. Cpa main core comprises the CnaB(M) domain (green), in whose fold the TED(T) domain is inserted (yellow). The TED(T) domain contains a thioester bond formed between the residues Cys426 and Gln575 (red), which mediates the attachment to cell-surface molecules. **b)** In the folded state, nucleophiles like methylamine (MA) can cleave the thioester bond and bind covalently to the Gln side chain (+MA); however, thioester bond reformation and ligand uncoupling (-MA) can occur. After mechanical extension, the presence (circle pathway) or absence (empty circle pathway) of the thioester bond can be assessed as a difference in the extension of the protein. **c)** Double-covalent magnetic tweezers experimental assay. Protein anchors SpyCatcher-HaloTag are covalently immobilized both to the surface of the glass and of the paramagnetic bead. A chimeric polyprotein made of four copies of Cpa and flanked by SpyTag peptides is covalently linked to the glass and to the bead through the reaction of the SpyCatcher/SpyTag split protein system. On the top of the scheme (not shown), the position of a pair of magnets is controlled for the application of calibrated forces to the tethered molecule. **d)** Magnetic tweezers recording of a Cpa polyprotein exposed to 100 mM methylamine, where the extension of the molecule is registered along time. A force pulse of 115 pN leads to the mechanical unfolding of the four Cpa domains, which is detected as stepwise increases in the extension. Here, three of the domains lack their internal thioester bond (empty circles) yielding an extension of ~95 nm, while one of the domains preserves its thioester bond (circle) and yields an unfolding extension of ~49 nm. Following a 100 s quench force pulse at 3 pN, which favors both folding and bond reformation, a second 115 pN pulse reveals that two Cpa domains reformed their thioester bonds (circles), decreasing the final extension of the polyprotein by 90 nm, as a consequence of the polypeptide sequence trapped by the newly formed bonds.

Here, we demonstrate a novel magnetic tweezers force spectroscopy approach to resolve in detail the force-dependency of Cpa thioester bond reactivity in the 3-115 pN force range. Unlike AFM, magnetic tweezers possesses an incomparable stability that grants access to days-long recordings on the same molecule, with millisecond and sub-pN resolution^17,18^. Cpa is a mechanically stable protein^11^, and, to apply high forces for long times, we designed a novel double-covalent anchoring strategy based on HaloTag chemistry and SpyCatcher/SpyTag split protein technique^19–21^, which allows for the end-to-end covalent immobilization of single Cpa polyprotein molecules. This technical advance enables us to explore different conditions on the same molecule without probe detachment, a limiting factor in force spectroscopy experiments^22–25^. With these improvements, we now determine the force-dependency of Cpa folding and its relation to thioester bond cleavage by the nucleophile methylamine. We find that methylamine-induced cleavage is inhibited at forces >35 pN, while thioester reformation and ligand uncoupling occur at forces <6 pN. Remarkably, protein folding precedes and is a requirement for thioester reformation, which suggests an allosteric role of folding on the reactivity of this bond. We hypothesize that the different force ranges over which thioester cleavage, reformation, and Cpa folding occur provide the bacteria with a novel mechanism to respond to varying levels of shear stress. At high shear stress spikes, like those that the bacterium might experience during a cough episode, the thioester bonds become unreactive and “lock up” above 35 pN with covalent-strength anchoring that can withstand forces over 1000 pN. When the shear stress eases up, the folding of the Cpa parent protein, at 6 pN or less, permits thioester bond reactivity and allows it to cleave and reform repeatedly, which permits a form of nomadic migration of the infecting pathogen. We dub such folding-controlled covalent reactivity: “smart covalent bond”. In the current context of antibiotic resistance^26^, targeting the bacterial adhesion molecules stands out as a promising strategy to battle infections^27^, especially considering the difficulties for treating those caused by Gram positive pathogens^28^. In such effort, we identify a mechanism for the abrogation of Cpa thioester bond reactivity towards surface ligands, based on the oxidation of the side chain thiol of the Cys residue involved in the thioester bond. A better understanding of the adhesive chemistries of Gram positive pathogens will permit the rational development of novel classes of antibiotics and vaccines, of great significance to society.

## Results

### Double-covalent magnetic tweezers anchoring

To explore Cpa thioester bond mechano-chemistry, we use a tetramer of the domains CnaBD595A(M)-TED(T) of this adhesin (**Figure 1a and b**), as we previously described^11^. The Cys426-Gln575 thioester bond resides within the TED domain, whose fold is contained inside the fold of the CnaB domain (**Figure 1a**). This protein exhibits high mechanical stability, and requires the application of high forces for unfolding. To solve this problem, we develop a strategy to covalently anchor Cpa polyproteins both to the glass surface and the magnetic probes of a magnetic tweezers setup. Both glass and probe surfaces are functionalized with the HaloTag ligand, which permits the covalent immobilization of HaloTag proteins^17,19,20^. First, we immobilize the chimeric protein SpyCatcher-HaloTag on both surfaces. Then, we add the chimeric polyprotein SpyTag-(CnaBD595A-TED)4-SpyTag to the glass surface, allowing the reaction with the SpyCatcher-HaloTag present on the surface. The SpyCatcher/SpyTag split protein system reacts to form an intermolecular isopeptide bond between the SpyTag and the SpyCatcher counterpart^21,29,30^, covalently connecting both chimeric proteins. Finally, we close this assembly by adding functionalized paramagnetic beads, whose surface-bound SpyCatcher-HaloTag protein reacts with the SpyTag peptide present on the free end of the Cpa polyprotein (**Figure 1c**). After capping, the Cpa polyprotein becomes covalently tethered both to the glass and bead surfaces.

The magnetic tweezers experiment starts when the protein-bound paramagnetic bead is exposed to a magnetic field^17^. The presence or absence of the thioester bonds in the Cpa polyprotein can be easily detected as a difference in the unfolding extensions (**Figure 1b**). **Figure 1d** shows a magnetic tweezers trajectory of a Cpa polyprotein which has been previously exposed to a solution containing 100 mM methylamine (Hepes 50 mM pH 8.5, NaCl 150 mM, ascorbic acid 10 mM, EDTA 1 mM). The application of a constant force of 115 pN leads to the sequential unfolding of the Cpa polyprotein, yielding stepwise increases in length of different sizes: one corresponding to thioester bond-intact proteins (~49 nm), and three corresponding to thioester bond-cleaved proteins (~95 nm). Subsequently, the force was reduced to 3 pN to allow the folding of Cpa and also the reformation of the thioester bonds. A second 115 pN pulse reveals that two more Cpa domains reformed their bonds (~49 nm steps), and the total extension of the polyprotein decreases by 90 nm, since the formation of these two thioester bonds prevents the full extension of the protein. The different extensions of Cpa, depending on the presence or absence of its internal thioester bond, serve to clearly identify the status of the bond.

### Force-dependency of the thioester bond cleavage and reformation

The mechanical unfolding of the CnaB_D595A_-TED domains with the thioester bond intact remains limited to the polypeptide sequence not-trapped by the bond. This accounts for a total of 164 residues located before the Cys426 and after the Gln575, which corresponds to the ~49 nm steps observed on **Figure 1d**. In a nucleophile-free solution, the polyprotein unfolding at 115 pN reveals stepwise increases in length of 48.6 ± 4.5 nm (mean±SD), as it can be seen in the trajectory of the **Figure 2a**. In these unfolding extensions, the entire CnaB fold and a small region of the TED domain are stretched—TED fold spans from residues Ala393 to Gly579. Due to the exquisite force resolution of magnetic tweezers, we can explore not only the Cpa tetramer unfolding at high forces, but also the reversible process of folding at low forces (**Figure 2b**). As it can be seen on **Figure 2a**, a quench for 100 s at 6 pN and a subsequent pulling at 115 pN shows no evidence of protein folding (P_f_=0.0). On the contrary, holding the protein at 4 pN for the same amount time is enough to completely fold the thioester-intact Cpa tetramer (P_f_=1.0), while at 5.5 pN only half of the domains could fold (P_f_=0.5). In this manner, we determine the folding probability of the thioester-intact polyprotein, which shows a sharp transition from fully folded at 4.5 pN, to completely unfolded at 6.5 pN (**Figure 2c**). Therefore, Cpa mechanical unfolding in the absence of nucleophiles yields a homogeneous population of steps of ~49 nm, which confirms that over the explored range of forces the thioester bond remains inert.

**Figure 2.**
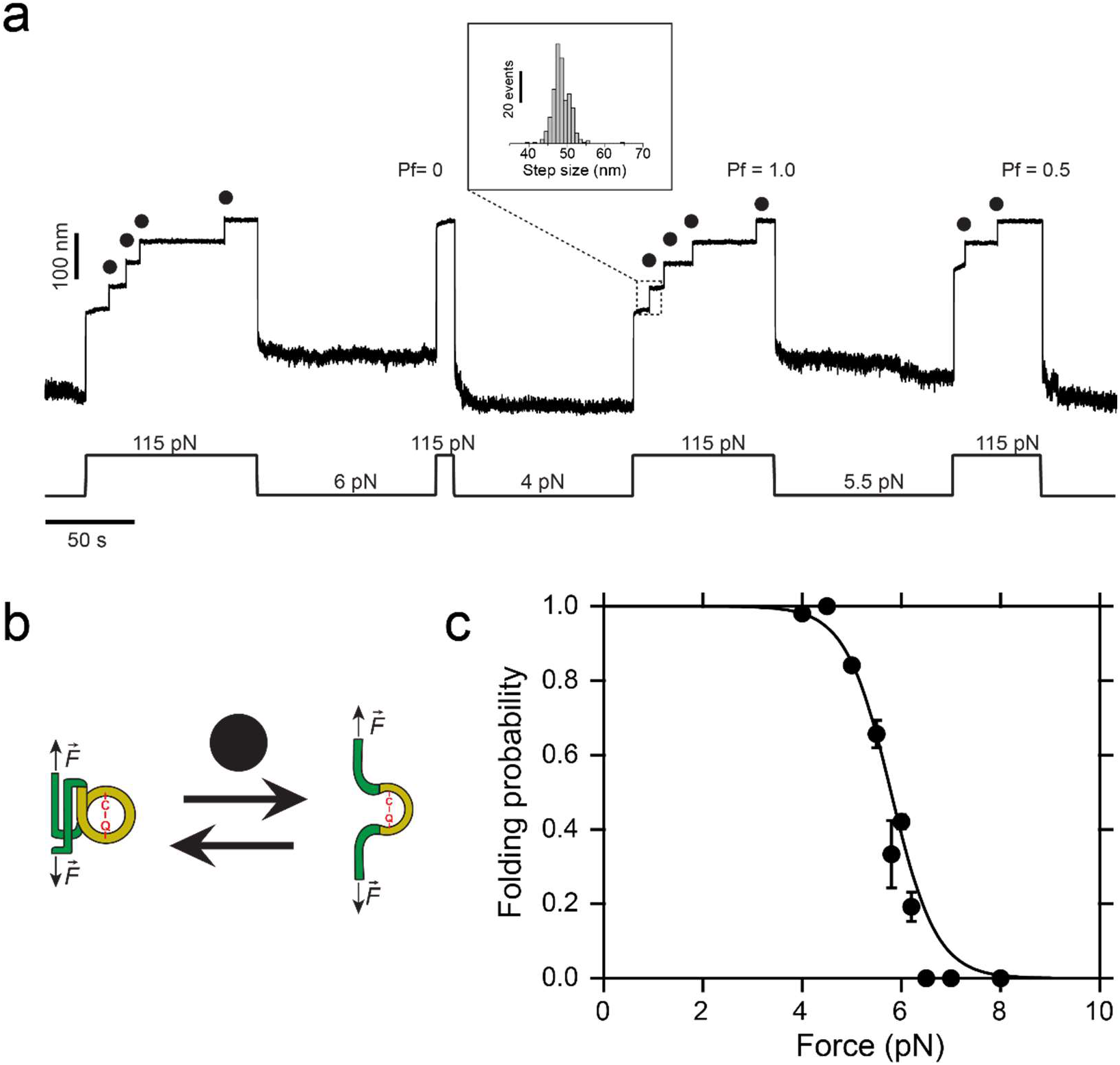
Dynamics of the thioester-intact Cpa polyprotein under force. **a)** Magnetic tweezers trajectory of the Cpa tetramer. High force pulses at 115 pN unfold the thioester-intact Cpa domains, which show 48.6 ± 4.5 nm (mean±SD, n=259) stepwise extensions (inset histogram). Low force pulses of 100 seconds long allow Cpa refolding, enabling us to determine the folding probability (Pf) at different forces. As an example, a quench at 6 pN does not allow folding of any of the domains, meanwhile the four fold at 4 pN (Pf=1.0), and only two fold at 5.5 pN (Pf=0.5). **b)** Cartoon representation of the folding-unfolding of the Cpa domain. The thioester bond between Cys426 and Gln575 clamps the TED domain (yellow), limiting its extensibility. **c)** Folding probability of thioester-intact Cpa. Data points are fitted to a sigmoidal function and they represent the calculated average folding probability and the error at each of the forces tested for 100 s, based on a jackknife estimator (n=49 at 4 pN; n=2 at 4.5 pN; n=12 at 5 pN; n=9 at 5.5 pN; n=4 at 5.8 pN; n=19 at 6 pN; n=8 at 6.2 pN; n=4 at 6.5 pN; n=8 at 7 pN; n=3 at 8 pN).

The stability of magnetic tweezers and the double-covalent anchoring of the protein allow us to exchange the solution in the experimental liquid chamber, enabling us to expose a single molecule to different conditions. Hence, to explore the thioester bond reactivity under force, we change to a solution containing 100 mM methylamine, which we add after the mechanical unfolding of the thioester-intact Cpa, as shown in **Figure 3a**. At 115 pN, the addition of methylamine does not yield any additional extension increase, indicating the lack of reactivity of the thioester bond at high forces. Taking advantage of the magnetic tweezers force resolution, we apply a protocol with consecutive decreasing force pulses of 100 s to elucidate the force-range reactivity of this bond in real time. Initially, decreasing the force to 30 pN does not alter the thioester bond state, as it can be seen from the following 115 pN pulse where the same final extension of the molecule is reached. By contrast, applying a pulse of 28 pN, reveals one discrete step originating from the bond cleavage of one of the four Cpa proteins. When we stretch again at 115 pN, the final extension of the molecule increases by 45 nm, which confirms this observation. This additional length comes from the release of the polypeptide sequence sequestered by the Cys426-Gln575, which scales with the number of residues previously trapped by the bond and also with the applied force following the freely-jointed chain model for polymer elasticity^31^ (**Supplementary Fig. 1**). Finally, dropping the force to 20 pN leads to the rapid cleavage of the three remaining bonds in the polyprotein, yielding three steps of 38 ± 3.2 nm (mean±SD, inset histogram #2). Exploring the range from 15 to 35 pN, we determine that thioester bond cleavage does not occur over a 100 s time-window if Cpa is exposed to forces >35 pN. When held at lower forces, stepwise increases in length occur due to thioester bond cleavage, reaching completion in 100 s at forces <23 pN (**Figure 3b**), which indicates a negative force-dependency in the ligand-induced cleavage. In **Figure 3c**, we show the rate of cleavage at four representative forces (20, 23, 25, and 27 pN), where in the case of 27 pN it can take up to >500 s to complete. The cleavage rate of the thioester bond decreases with the mechanical load, which underpins the negative force-dependency of this reaction. Assuming the Bell model for bond lifetimes under force^32^, this negative exponential trend indicates a negative distance to the transition state of ~8 Å, which suggests that thioester bond lysis requires a structural shortening of the protein conformation.

**Figure 3.**
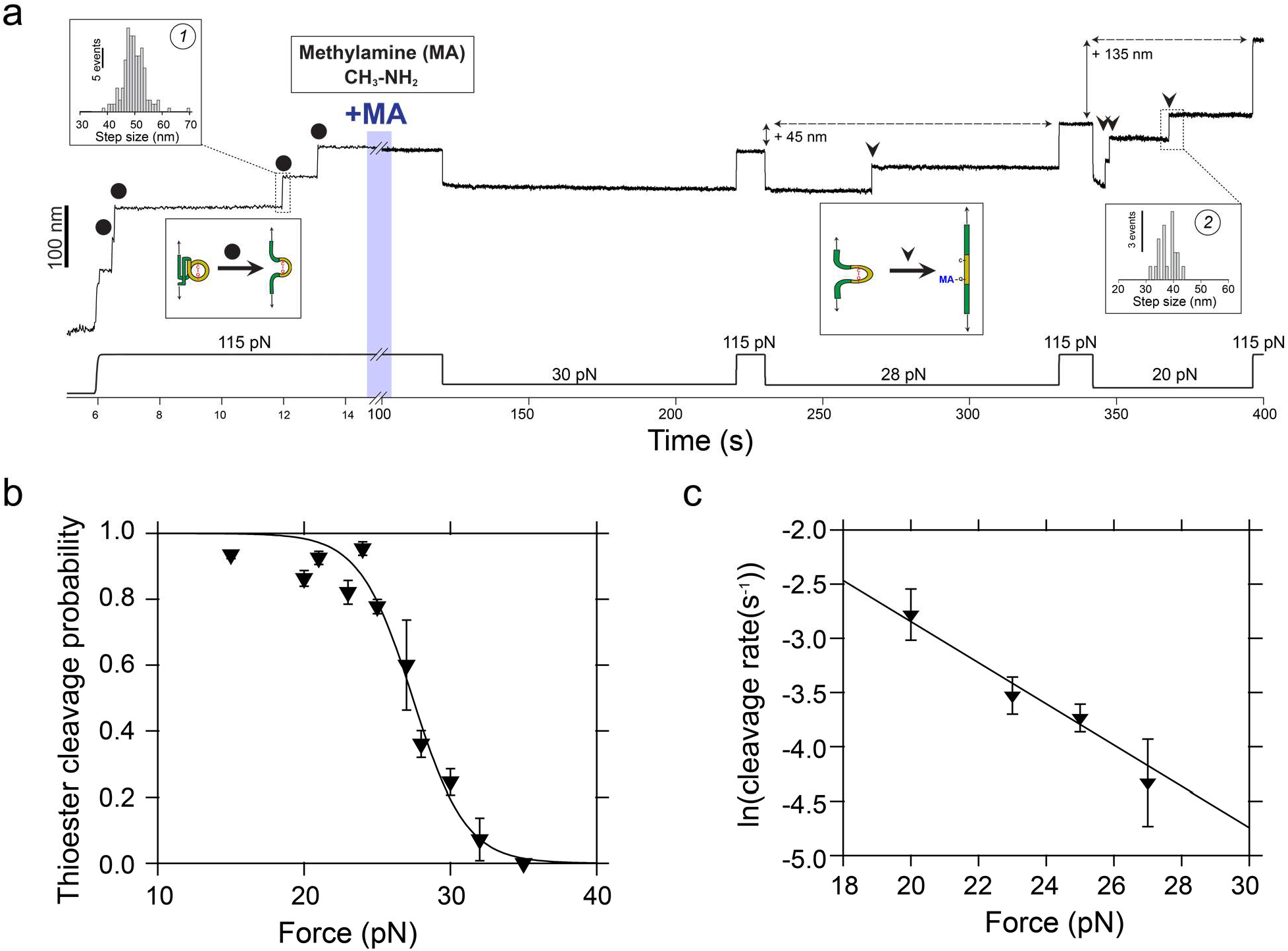
Cpa thioester bond cleavage is negatively force-dependent. **a)** Magnetic tweezers trajectory of the Cpa polyprotein. After the unfolding of the thioester-intact Cpa domains at 115 pN (circles; histogram inset #1: 49.4±4.5 nm, mean±SD, n=165), the buffer is exchanged to a Hepes solution containing 100 mM methylamine (+MA). At high force, no additional steps are registered as it would be expected from a thioester bond cleavage event. Thereafter, we apply a protocol with subsequent pulses of decreasing mechanical load to investigate the force-dependency of the reaction. While at 30 pN no cleavage is observed, 100 s at 28 pN reveal one step that comes from the methylamine-induced cleavage of the thioester bond of one of the four Cpa domains (triangle). At 115 pN, the final extension of the molecule has increased by 45 nm, which originates from the polypeptide sequence released after thioester bond lysis. When held at 20 pN, the three remaining thioester bonds are cleaved (triangles; histogram inset #2: 38±3.2 nm, mean±SD, n=20) and the final extension of the molecule increases for another 135 nm. **b)** Thioester bond cleavage probability as a function of force measured over a 100 s time-window. Data points show the average and the error based on a jackknife estimator. The line represents a sigmoidal fit to the data (n=15 at 15 pN; n=12 at 20 pN; n=8 at 21 pN; n=9 at 23 pN; n=7 at 24 pN; n=12 at 25 pN; n=5 at 27 pN; n=7 at 28 pN; n=10 at 30 pN; n=3 at 32 pN; n=4 at 35 pN). **c)** Rate of thioester bond cleavage as a function of force. Data points show the natural logarithm of the cleavage rate and the bars show the standard error of the mean. The line represents an exponential fit to the data, whose slope indicates a negative distance to the transition state x^†^=-0.78 nm, which suggests a requirement of a contraction of the Cpa polypeptide substrate to proceed with the cleavage of the bond, explaining its negative force-dependence (n=24 at 20 pN; n=21 at 23 pN; n=19 at 25 pN; n=21 at 27 pN). Only molecules where the cleavage reaction reached completion for all the candidate domains of the polyprotein were taken into account.

Our results indicate that thioester bond cleavage is hindered at forces >35 pN, and that the kinetics of this reaction are steeply affected by the mechanical load. Methylamine-induced cleavage leads to the covalent binding of this nucleophile to the Gln side chain, but the backwards reaction involving thioester reformation and ligand uncoupling can occur in the folded state of Cpa. To explore this opposite reaction, we design the force protocol shown on **Figure 4a**. After mechanical unfolding of the Cpa polyprotein, and the cleavage of the thioester bonds with methylamine (see **Figure 4b**), we wash the nucleophile out of the reaction buffer and reduce the force on the protein, to favor both the bond reformation and folding of the protein. These conditions allow us to observe a sharp increase in the reformation probability once the Cpa protein is exposed to forces <6 pN (**Figure 4c**). The number of reformation events—detected as thioester-intact Cpa unfolding steps at 115 pN—scales with the number of cleavage events observed before the methylamine washout. Interestingly, the bond reformation force range closely tracks that of the folding of thioester-intact Cpa proteins. Given that the Cys and the Gln residues are moved away after cleavage, the force must be decreased to bring close the Cys thiol to attack the Gln carbonyl group and reform the thioester. The fact that Cpa folding occurs at higher forces entails that folding precedes the thioester bond reformation, as it has been also described for the formation of disulfide bonds^33,34^.

**Figure 4.**
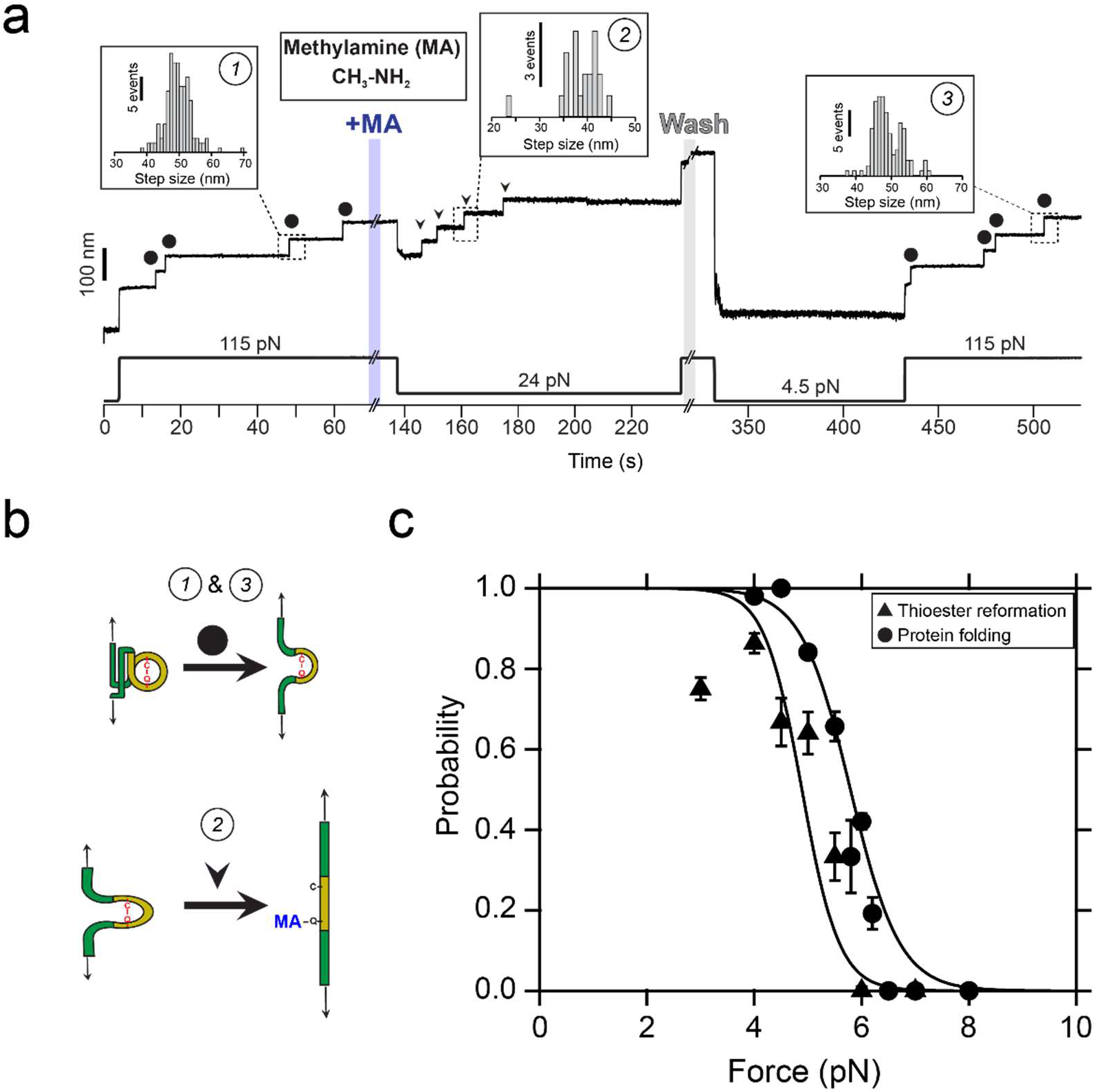
Protein folding drives thioester bond reformation. **a)** Magnetic tweezers trajectory of the Cpa polyprotein. After the unfolding of the thioester-intact Cpa domains at 115 pN (circles; inset histogram #1: 49.4 ± 4.5 nm, mean±SD, n=165), the buffer is exchanged and the polyprotein is exposed to a solution containing 100 mM methylamine (+MA). As expected, we do not observe cleavage at this high force, but a drop to 24 pN permits the full cleavage of the four candidate thioester bonds (arrows, inset histogram #2; 38.4 ± 4.3 nm for 24 pN, mean±SD, n=21). To study the reformation of the bond, we remove the nucleophile-containing buffer at high force, and quench the force to 4.5 pN for 100 s to favor bond reformation and protein folding. We stretch again the polyprotein at 115 pN and identify four thioester-intact Cpa domains, which indicates that the four cleaved candidates were able to fold and to reform their bonds (circles; inset histogram #3: 48.8 ± 4.1 nm, mean±SD, n=117). **b)** Cartoon representation of the extension events registered on the Cpa trajectory shown in **a)**. Events #1 and #3 show the mechanical extension at 115 pN of thioester-intact Cpa, before cleavage and after reformation, respectively. Event #2 shows the extension after methylamine (MA) cleavage at 24 pN. **c)** Comparison between the thioester bond reformation (upwards triangles and sigmoidal fit) and the thioester-intact Cpa folding probability (circles and sigmoidal fit, from **Figure 2c**) as a function of the mechanical load. Data points for reformation show the average and the error based on a jackknife estimator. Reformation registered as the amount of thioester-intact domains after methylamine washout and after a 100 s time-window at the folding/reformation force range (n=12 at 3 pN; n=11 at 4 pN; n=7 at 4.5 pN; n=7 at 5 pN; n=5 at 5.5 pN; n=4 at 6 pN; n=3 at 7 pN).

### Blocking the thioester bond reformation

Our experiments with methylamine demonstrate the full reversibility of the cleavage reaction when Cpa is held at low forces and allowed to fold. These experimental conditions resemble the kind of interactions that Cpa adhesin could establish with the host ligands, binding and unbinding depending on the mechanical load experienced at the bond interface. From a therapeutic perspective, the irreversible thioester bond cleavage by a ligand analog would prevent bacterial adhesion, easing the bacterial removal from the tissues by the host’s clearance mechanisms— mucus flow, coughing, etc. Taking into account the Cys residue side chain, we explored the cleavage and reformation of Cpa thioester bond after the treatment with cystamine, another primary-amine nucleophile which contains a disulfide bond in its structure. Following the same protocol as with methylamine, we first unfold thioester-intact Cpa proteins (**Figure 5a**, inset histogram #1) and then introduce a solution containing 100 mM cystamine. Upon force reduction to 25 and 20 pN for 100 s, the cleavage steps appear as it occurred with methylamine (inset histogram #2). After cystamine removal from the solution, the protein is allowed to refold and to reform the thioester bonds at 4 pN. If bond reformation occurs, at 115 pN we should detect the same ~49 nm steps registered before cystamine treatment. However, 95 ± 10.7 nm single steps (mean±SD, inset histogram #3) appear, which account for the full extension of thioester-cleaved Cpa proteins. Despite attempts to reform the bonds by reducing the force for several cycles (**Supplementary Figure 2**), we can only detect full Cpa unfolding steps after cystamine. This nucleophile’s disulfide bond can be attacked by Cys426 free thiol to generate an intermolecular disulfide bond (diagram on **Figure 5b**). Cys426 thiol oxidation would prevent thioester reformation, which could explain our observations where bond reformation is never observed after cystamine intervention. To further test this hypothesis, we add the reducing agent TCEP, to reduce disulfide bonds and liberate the Cys426 thiol. After solution exchange and force reduction, we observe again at high force the unfolding steps of thioester-intact Cpa proteins (**Supplementary Figure 3**). **Figure 5c** compares the cleavage and the reformation probability of the thioester bond after treatment with methylamine and cystamine. While both nucleophiles exhibit the same cleavage behavior at 20 and 25 pN, reformation at 4 pN is completely abolished after cystamine treatment. However, if cystamine-blocked proteins are treated with a solution containing 10 mM TCEP, the thioester bond recovery reaches the same values as with methylamine. These findings strongly support the idea that cystamine blocking activity relies on the formation of an intermolecular disulfide bond with Cpa Cys426 side chain, which prevents thioester bond reformation and which can only be rescued after the action of a reducing agent.

**Figure 5.**
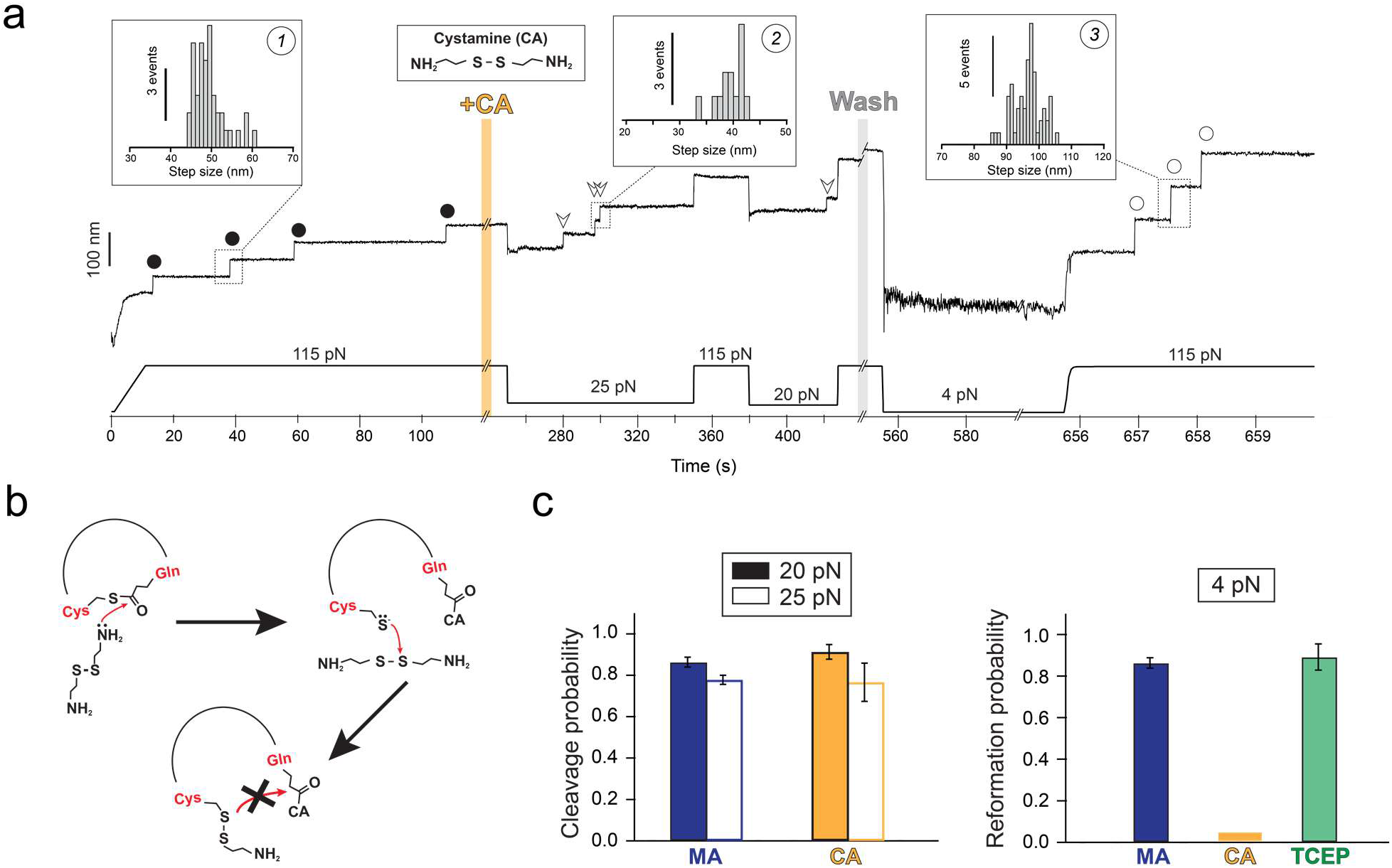
Cystamine-mediated abrogation of Cpa thioester bond reformation. **a)** Magnetic tweezers trajectory of the Cpa polyprotein. After the unfolding of the thioester-intact Cpa domains at 115 pN (circles, inset histogram #1; 49.3 ± 3.8 nm, mean±SD, n=42), the buffer is exchanged and the polyprotein is exposed to a solution containing 100 mM cystamine (+CA). At 115 pN, no additional extensions are registered, but a drop in the force to 25 pN for 100 s leads to the appearance of three steps which account for the release of the polypeptide sequence trapped by the thioester bonds (empty arrows, inset histogram #2; 39.3 ± 2.4 nm for 25 pN, mean±SD, n=13). After the cleavage of all the bonds and after cystamine washout, force is quenched to 4 pN for 100 s to favor folding and reformation of the thioester. The final 115 pN pulse reveals three steps corresponding to thioester bond-cleaved Cpa domains (empty circles, inset histogram #3; 95 ± 10.9 nm, mean±SD, n=58). **b)** Chemical scheme depicting the reformation blocking effect of cystamine (CA). After the thioester bond nucleophilic cleavage by one of the CA primary amines, the free Cys thiol can attack CA disulfide bond (from the bound CA, or from another CA molecule). As a result, an intermolecular disulfide bond between Cpa Cys426 and CA is formed, preventing the thioester bond reformation. This disulfide reshuffling breaks the CA molecule and generates one free cysteamine molecule (not shown in the scheme), and a Cys426-bound cysteamine. **c)** Left panel compares the thioester bond cleavage probability by methylamine (MA) and cystamine (CA) at 20 and at 25 pN (MA, n=12 at 20 pN, n=12 at 25 pN; CA, n=8 at 20 pN, n=5 at 25 pN). Right panel compares the thioester bond reformation probability after 100 s at 4 pN after the treatment with MA, CA, and after the treatment with CA followed by TCEP (MA, n=11; CA, n=11; TCEP, n=5).

## Discussion

Bacteria pathogens possess molecular traits that enable host colonization under mechanical stress. Among these, isopeptide bonds stand out by conferring high mechanical and thermal stability to the adhesive proteins and pili of Gram positive bacteria^35–37^. These bonds preserve the mechanical integrity of the bacterial anchors^38–40^, but ultimately the adhesion lifetime relies on the properties and the strength of the bacteria receptor-host ligand interaction. Gram positive adhesin-ligand binding has evolved to withstand nanoNewton-scale mechanical loads, like *Staphylococcus epidermidis* SdrG adhesin^41^, but also to respond to force in a putative catch bond-like manner, such as *Staphylococcus aureus* ClfA and Clfb adhesins^42,43^, or *Streptococcus pneumoniae* pilin RrgB^44^ In the catch bond mechanism, force triggers conformational changes on the adhesin structure that increase the bond lifetime with the ligand, enabling the bacteria to respond to force thresholds^45^. Most of these adhesins interact with extracellular matrix proteins—such as fibrinogen and collagen^46,47^—and establish non-covalent bonds with their ligands. In addition to these, it was recently discovered the existence of thioester bond-adhesins in some Gram positive organisms^5–7^. These adhesins can form a covalent bond with the substrate through the nucleophilic attack of its thioester bond by a primary-amine ligand, like the ε-amino group of a Lys residue^5^. Nevertheless, the establishment of an irreversible covalent anchoring would impose a sessile strategy on the cell, hindering its spreading and colonization^48^. Experiments with *S. pyogenes* Cpa adhesin revealed that in the absence of force the thioester bond cleavage by soluble nucleophiles and its reformation existed in equilibrium; however, the application of tensile stress to the thioester bond prevented both its cleavage and reformation, indicating that force modulates the reactivity of this bond^11^. Intramolecular thioester bonds are uncommon in the structure of proteins, having been only identified in the immune complement proteins, in α2-macroglobulin anti-protease^49–51^, and in Gram positive adhesins^6,7^. In the case of non-activated complement proteins, nucleophilic cleavage and reformation can occur^52^, but the proteolytic activation of these proteins leads to a rapid and irreversible binding to its target substrates^53^, which contrasts with the reversible and force-modulated reactivity of *S. pyogenes* adhesin.

Here, using magnetic tweezers force spectroscopy and a new protocol for the covalent anchoring and assembly of polyproteins, we identify the force range for Cpa thioester bond reactivity. Our results indicate that ligand-induced cleavage occurs at forces <35 pN. This impaired reactivity under force contrasts with the positive effect of force on the mechano-chemical cleavage of disulfide bonds by small reducing agents. These disulfide reductions proceed via an S_N_2 mechanism that experiences a ~0.3 - 0.4 Å elongation to the transition state^13,14^. On the contrary, enzymatically-catalyzed disulfide reductions by thioredoxin exhibit a negative force-dependency, where the substrate polypeptide must contract under force in order to align with key catalytic residues of the enzyme^15,54,55^. A similar mechanism of polypeptide contraction may underlie the observed force-dependency of the Cpa thioester bond, as it can be inferred from the negative distance to the transition state we have observed. We explain the negative force-dependency of thioester cleavage as an autocatalytic mechanism that facilitates the nucleophilic attack, as it has been reported in a close Cpa homolog in *S. pyogenes*^8^; mechanical unfolding would disrupt the thioester active site and inhibit bond cleavage, deforming the spatial arrangement of key catalytic residues placed in the vicinity of the Cys-Gln bond. Notably, we observe small stepwise fluctuations at forces below 35 pN, which precede thioester cleavage and disappear after the reaction occurs (**Supplementary Figure 4**). However, no single discrete step size population is apparent and we cannot assign a specific structural transition to these fluctuations. Nevertheless, the close temporal relationship of these fluctuations to thioester cleavage events suggests a requirement of some structural contraction, and in turn a negative force-dependency to the reaction rate. In the backward reaction, the Cys426 and Gln575 residues must be in close proximity for thioester reformation, which is most probable at or close to the native folded state. Supporting this mechanism, we measured the folding force-dependency of thioester-intact Cpa (**Figure 2c**), which closely tracks the profile of thioester reformation (**Figure 4c**). This pathway of refolding followed by reformation can explain the sharp transition in the force-dependency from 4 pN to 6 pN, with reformation restricted to the force range of protein folding. This behavior shows an analogy with the process of enzymatic-assisted oxidative folding, where protein folding brings in close proximity the Cys residues involved before disulfide bond formation can proceed^33,56^.

Significantly, our experimental pulling axis from the N and C ends of Cpa is not the physiological one, although simulation of the *in vivo* pulling axis between the C-terminus and Gln575 reveals mechanical deformation of the thioester active site^11^. Indeed, subsequent experimental work with the proper pulling geometry will be needed to refine the force range of this transition. In any case, our findings about the force-dependency of Cpa thioester bond reactivity allow us to hypothesize that the bond reactivity modulation by protein folding can be used as a smart binding solution in the Gram positive adhesion strategy, which simultaneously solves the mobility and the anchoring problem (**Figure 6**). Non-covalent catch bonds show an increased lifetime at certain levels of mechanical load, but exceeding forces terminate the binding^45^. Thioester bonds would operate in the low force regime (<6 pN) as catch bonds, where surface ligands cleave them but reformation can occur, as long as the protein remains in the native folded state. Under these conditions, soluble ligands like histamine—which is released at infection sites^57^—can also bind to Cpa and compete with the surface targets of the adhesin. However, the lack of tensile stress in the Gln575-histamine interface would allow the Cys-Gln thioester bond reformation, which would release the histamine and reset Cpa ready for another incoming ligand. By contrast, increasing mechanical loads on those bonds established with surface-bound ligands at low force would prevent reformation due to the mechanical deformation and unfolding of Cpa—mechanical allostery—, inducing a long-lived covalent bond able to survive nanoNewton-scale perturbations^58^. Only after the mechanical challenge is finished and the force is reduced, Cpa folding and bond reformation can occur to favor cell rolling again. Hence, this folding-modulated reactivity of thioester bonds places them as smart covalent bonds, which have the ability to provide the bacteria with a balanced strategy: to switch from a nomadic mobility phase at low shear stress that optimizes cell spreading, to a locked phase under harsh mechanical conditions that induce dislodgement.

**Figure 6.**
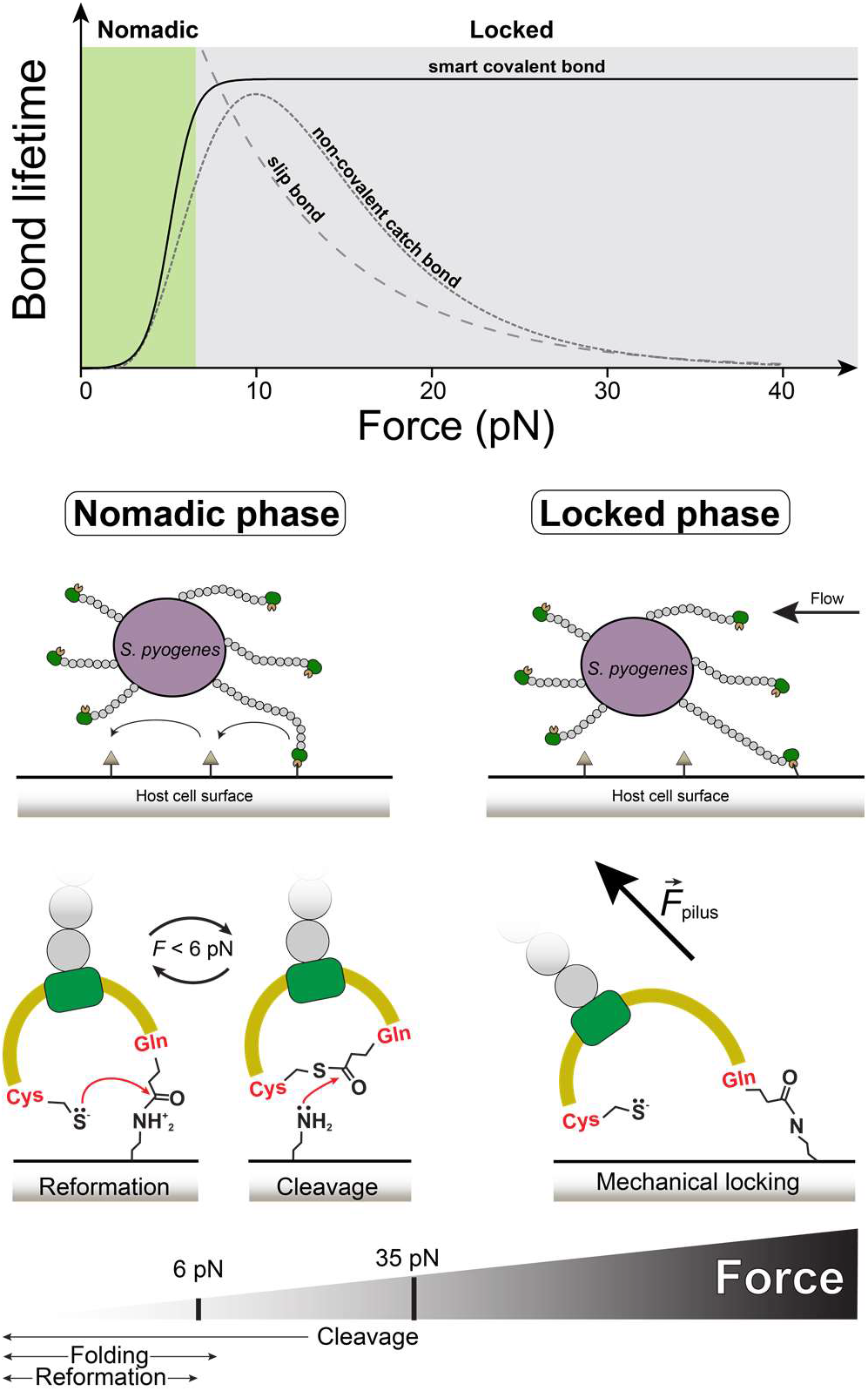
Cpa thioester bond allosteric modulation by folding enables the bacterium to switch the mobility strategy depending on the shear stress. Top graph compares bond lifetimes as a function of the mechanical load for slip bonds, non-covalent catch bonds, and thioester smart covalent bonds (slip bond and catch-bond data adapted from^59^, plotted in arbitrary units). While increasing loads exponentially decrease the lifetime in slip bonds, in non-covalent catch-bonds it increases; however, loads above certain threshold decrease the lifetime. The adhesin thioester bond is allosterically modulated by force, establishing shortlived bonds with its ligand substrates at low mechanical stress when the protein is folded, but turning into a long-lived bond that permits the bacterium to remain attached under large mechanical challenges. We hypothesize that these smart covalent bonds allow bacteria to switch between a nomadic mobility phase at low force—where cleavage and reformation exist in equilibrium at forces <6 pN in the native folded state of the protein, and the cell rolling over surfaces is favored—to a mechanically locked phase at larger loads, which would unfold and deform the Cpa-bound protein structure, preventing Cys426 side chain thiol to attack back the Gln575 side chain and reform the thioester bond.

Our results indicate that protein folding can modulate the binding activity of *S. pyogenes* Cpa, and they also indicate that chemically targeting the Cys-Gln thioester bond can be of potential interest for the development of antiadhesive drugs. The inhibitory effect observed after the treatment with cystamine indicates that, after nucleophilic cleavage, disulfide bond exchange occurs between Cys426 free thiol and cystamine disulfide bond, arresting the reformation. This conclusion is supported by the regenerative effect registered after TCEP treatment, which reduces the cystamine-Cpa intermolecular disulfide bond and frees the Cys426 thiol, enabling the reformation reaction. While methylamine and histamine transiently cleave the thioester bond, bifunctional soluble ligands with nucleophilic and thiol oxidation activities could permanently bind to Cpa to disable its adhesin function, establishing a new therapeutic path to tackle the antibiotic-resistance problem.

## Methods

For additional methods, see the Supplementary Information.

### Protein purification

All the reagents employed in this research were from Sigma-Aldrich, unless otherwise specified. The SpyTag-(Cpa)_4_-SpyTag and the HaloTag-SpyCatcher constructs were cloned into the expression plasmid pQE80L (Qiagen). Protein expression and purification was done as described elsewhere^36^. In brief, *E. coli* ERL cells were transformed with the pQE80L-SpyTag-(Cpa)_4_-SpyTag plasmid or pQE80L-SpyCatcher-HaloTag, and protein expression was induced with 1 mM Isopropyl β-D-1-thiogalactopyranoside overnight at 25°C or 37°C, respectively. Cells were lysed in a French press (Sim-Aminco), and proteins were purified from the lysate with the His60 Ni Superflow Resin (Clontech). An additional purification step was done through size exclusion chromatography in a Superdex 200 FPLC column (GE Healthcare).

### Bead surface functionalization

10^8^ amine coated Dynabeads M270 (Thermo Fisher Scientific) were washed in PBS buffer pH 7.4 and incubated in a PBS solution containing 1 % v/v glutaraldehyde for 1 h in a rotator at 18 rpm (Labnet). After extensive washing, the beads were incubated in a PBS solution containing 25 μg/mL of the HaloTag ligand O4 (Promega) for at least 4 h at constant rotation (room temperature). After washing, beads were treated with blocking buffer (Tris-HCl pH 7.4, NaCl 150 mM, NaN_3_ 0.001% w/v, and 1% w/v of sulfhydryl-blocked BSA (Lee Biosolutions)), overnight at 4°C and at constant rotation. Optimal bead protein functionalization was achieved with a 15:5 μM ratio of HaloTag protein (also expressed and purified using the method explained above) and SpyCatcher-HaloTag, respectively, for at least 12 hours at 4°C and at constant rotation.

### Fluid chamber functionalization

Magnetic tweezers experiments were conducted on fluid chambers made of two sandwiched glasses (Ted Pella) of 24×40 mm (bottom) and 22×22 (top) separated by a thin parafilm template. Prior to fluid chamber assembly, bottom glasses were silanized with an ethanol solution containing 0.1 % v/v of (3-aminopropyl)-trimethoxysilane, and the top glasses were treated with repel silane for hydrophobization. Once assembled, the fluid chambers were exposed to a PBS pH 7.4 solution containing 1% glutaraldehyde to subsequently immobilize the reference beads (Spherotec) and the HaloTag ligand (Promega).

### Double covalent and molecular assembly

Fluid chambers were incubated with 5 μM SpyCatcher-HaloTag for 30 minutes. After an extensive rinse with Hepes buffer, the chambers were incubated with 5 μM SpyTag-(CnaB-TED)_4_-SpyTag for at least 1 h, and then extensively rinsed again. Once the fluid chamber was placed on the microscope, 20 μL of a 1:10 dilution of HaloTag:SpyCatcher-HaloTag functionalized beads were added to the fluid chamber and recirculated twice. Then, beads were allowed to react with the surface-bound molecules for 5 minutes before approaching the magnets and starting the experiment.

### Magnetic tweezers force spectroscopy

Force spectroscopy experiments were conducted on a custom-built magnetic tweezers apparatus, as previously described^17^. All the experiments were started applying a force of 4 pN. The unfolding pulses were done at 115 pN, until the complete unfolding of the thioester-intact Cpa domains (~49 nm steps). Only molecules showing the initial unfolding of 3 or 4 domains were considered. Buffer exchange to add or remove nucleophile molecules was done at 115 pN. Upon nucleophile addition to the fluid chamber, thioester bond cleavage was monitored over a 100 s time-window at forces ranging from 15 to 35 pN. Then, a 115 pN pulse was applied to monitor and compare the final extension of the molecule before and after the nucleophile treatment. At this high force, the nucleophile-containing buffer is washed out and then the force is quenched for 100 s at forces ranging from 3 to 7 pN, to favor refolding and thioester bond reformation. After, the folding and the thioester bond status of the domains are evaluated with a 115 pN pulse. The buffer used along the experiments contained 50 mM Hepes pH 8.5, 150 mM NaCl, 1 mM EDTA, 10 mM L-ascorbic acid, and was supplemented with 100 mM methylamine or cystamine for the thioester bond cleavage. To induce thioester bond reformation after cystamine treatment, the same buffer but supplemented with 10 mM TCEP was added and the force quenched to 4 pN to favor folding and reformation.

### Analysis

Analysis was done with Igor Pro 8.0 software (Wavemetrics). Recordings were smoothed using a 4^th^ order Savitzky-Golay filter with a box size of 51 points. Folding, cleavage, and reformation probabilities were calculated using a jackknife estimator for the calculation of the average probability and the standard deviation.

## Acknowledgments

This research was supported by the National Institutes of Health grant GG014033 (J.M.F). A.A-C. and R.T-R. express their gratitude to Fundación Ramón Areces (Madrid, Spain) for the financial support. We thank Carmen L. Badilla for reading and reviewing the manuscript.

## Author contributions

D.J.E., A.A-C. and J.M.F designed the research. A.A-C, D.J.E., R.T-R., S.H., E.C.E carried out the experiments. A.A-C, D.J.E., R.T-R. analyzed the data. A.A-C., D.J.E. and J.M.F. wrote the manuscript.

