## Supplemental Information for "Protein folding modulates the adhesion strategy of Gram positive pathogens"

### SUPPLEMENTARY INFORMATION

### Supplementary Methods

#### Protein engineering and expression

All the reagents employed in this research were from Sigma-Aldrich, unless otherwise specified. *S. pyogenes* Cpa gene was kindly provided by Mark Banfield (John Innes Centre, Norwich, UK). The gene was modified to include a 5'-end BamHI restriction site, a point mutation D595A to abolish CnaB(M) intramolecular isopeptide bond formation, and 3'-end BglII and KpnI restriction sites, as described previously[1]. A polyprotein containing four copies of Cpa—CnaB(M)-TED(T)—was assembled through successive cloning steps involving BamHI, BglII, and KpnI restriction sites, using pT7Blue (Novagen) as the cloning plasmid. The construct was then digested with BamHI/BglII and cloned into the expression plasmid pQE80L (Qiagen), which carries a N-terminal His tag. This plasmid was previously modified to contain two copies of the SpyTag sequence with a BamHI restriction site in between, which was digested to allow the insertion of the construct, generating the SpyTag-$(Cpa)_4$-SpyTag construct. All the cloning and amplification steps were done in XL10-Gold *E. coli* cells (Agilent Technologies). The probe and surface anchor protein, SpyCatcher-HaloTag, was also assembled using this same approach, and finally transferred to the pQE80L expression plasmid. The C-terminal HaloTag protein version was used for this construct[2].

Protein expression and purification was done as described elsewhere[3]. In brief, *E. coli* ERL cells (kindly provided by R.T. Sauer from Massachusetts Institute of Technology) were transformed with the pQE80L-SpyTag-$(Cpa)_4$-SpyTag plasmid or pQE80L-SpyCatcher-HaloTag, and protein expression was induced with 1 mM Isopropyl β-D-1-thiogalactopyranoside overnight at 25°C or 37°C, respectively. Cells were lysed in a French press (Sim-Aminco), and then the proteins were purified from the lysate with the His60 Ni Superflow Resin (Clontech). An additional purification step was done through size exclusion chromatography in a Superdex 200 FPLC column (GE Healthcare), eluting the proteins in 10 mM Hepes (pH 7.2), 150 mM NaCl, 1 mM EDTA (Hepes buffer). In the case of SpyCatcher-HaloTag protein, Hepes buffer additionally contained 10% v/v of glycerol. Purified proteins were aliquoted and frozen at -20°C until their use.

**Bead surface functionalization**

$10^8$ amine coated Dynabeads M270 (Thermo Fisher Scientific) were washed in PBS buffer, pH 7.4 and incubated in a PBS solution containing 1% v/v glutaraldehyde for 1 h in a rotator at 18 rpm (Labnet). After extensive washing, the beads were incubated in a PBS solution containing 25 µg/mL of the HaloTag ligand O4 (Promega) for at least 4 h at constant rotation. After washing, beads were treated with blocking buffer, which contains Tris-HCl pH 7.4, NaCl 150 mM, $NaN_3$ 0.001% w/v, and 1% w/v of sulfhydryl-blocked BSA (Lee Biosolutions), overnight at 4°C and at constant rotation. Optimal bead protein functionalization was achieved with a 15:5 µM ratio of HaloTag protein and SpyCatcher-HaloTag, respectively, for at least 12 hours at 4°C and at constant rotation. This 3:1 proportion results in an optimal bead surface coverage that prevents the formation of multiple tethers, since only the SpyCatcher-HaloTag molecules will serve as anchors for the glass surface-bound proteins. Beads were stored under this condition until use, moment in which they were extensively washed to remove unbound protein.

**Fluid chamber functionalization**

Magnetic tweezers experiments were conducted on fluid chambers made of two sandwiched glasses (Ted Pella) of 24x40 mm (bottom) and 22x22 (top) separated by a thin parafilm template trimmed with a laser cutting machine (Superland). The templates have a bow tie-like shape that allows the immobilization of the top glass over the bottom glass, and the formation of one well on each end of the bottom glass, which permits the exchange of buffer along the experiments. Prior to fluid chamber assembly, bottom glasses were washed and sonicated for 20 minutes in Hellmanex 1% (Helma), acetone, and ethanol. After the wash, the glasses were dried and exposed to air plasma for 15 minutes. Then, glasses were silanized for 20 minutes with an ethanol solution containing 0.1 % v/v of (3-aminopropyl)-trimethoxysilane, followed by several washes in ethanol. Finally, the glasses were dried with air, baked at 100°C for more than 20 minutes, and stored in a desiccator until further use. Top glasses were sonicated for 20 minutes in Hellmanex 1%, washed with ethanol, dried with air, and dried at 100°C for 10 minutes. Then, the top glasses were treated with repel silane for 30 minutes to make them hydrophobic. After, the glasses were dried with air, baked for 20 minutes at 100°C, and stored in a desiccator until use.

Fluid chambers assembly was done sandwiching the parafilm bow tie templates between the bottom and the top glasses, placed over a hot plate at 85°C, and with a flat 1 kg aluminum block pressing it. After 10 minutes, the fluid chambers were removed from the plate and a solution of PBS pH 7.4 with glutaraldehyde 1% v/v was flowed into the chambers and let to react for 1 h. After, a PBS solution containing 0.02 % w/v of 3.5-3.9 µm amine-coated polystyrene beads (Spherotech) was flowed and incubated for 20 minutes. After washing extensively, a PBS solution containing 25 µg/mL of the HaloTag ligand O4 was incubated overnight at room temperature. Finally, the fluid chambers were washed, blocked with blocking buffer overnight at room temperature, and stored at 4°C until further use.

**Double covalent and molecular assembly**

Fluid chambers were incubated with 5 µM SpyCatcher-HaloTag for 30 minutes. After an extensive rinse with Hepes buffer, the chambers were incubated with 5 µM SpyTag-(CnaB-TED)4-SpyTag for at least 1 h, and then extensively rinsed again. Once the fluid chamber was placed on the microscope, 20 µL of a 1:10 dilution of HaloTag:SpyCatcher-HaloTag functionalized beads were added to the fluid chamber and recirculated twice. Then, beads were allowed to react with the surface-bound molecules for 5 minutes before approaching the magnets and starting the experiment.

**Magnetic tweezers force spectroscopy**

Force spectroscopy experiments were conducted on a custom-built magnetic tweezers apparatus, as previously described[4]. The experimental fluid chambers are placed on the top of an inverted microscope (Olympus IX-71/Zeiss Axiovert S100) and illuminated with a collimated cold white LED (ThorLabs). The reference beads and the protein-bound paramagnetic beads are visualized employing a 100X oil-immersion objective (Zeiss/Olympus), which is mounted on a nanofocusing piezo actuator (P-725; Physik Instrumente). Image acquisition was done using a CMOS Ximea MQ013MG-ON camera, and image processing was done with a custom-written C++/Qt software. Data acquisition and piezo position control were done using a multifunction DAQ card (NI USB-6289, National Instruments). Proteins were exposed to calibrated forces using a pair of magnets mounted on the top of a voice-coil (Equipment Solutions) placed above the experimental fluid chamber. Magnets position was maintained under electronic feedback with a PID controller.

**Single molecule magnetic tweezers experiments on thioester bond cleavage and reformation**

All the experiments were started applying a force of 4 pN, which lifts the protein-bound beads from the surface and prevents nonspecific interactions. The unfolding pulses were done at 115 pN, until the complete unfolding of the thioester-intact Cpa domains (~49 nm steps). Only molecules showing the initial unfolding of 3 or 4 domains were considered. Buffer exchange to add or remove nucleophile molecules was done at 115 pN. Upon nucleophile addition to the fluid chamber, thioester bond cleavage was monitored on 100 s time windows at forces ranging from 15 to 35 pN. Then, a 115 pN pulse was applied to monitor and compare the final extension of the molecule before and after the nucleophile treatment. At this high force, the nucleophile-containing buffer is washed out and then the force is quenched for 100 s at forces ranging from 3 to 7 pN, to favor refolding and thioester bond reformation. After, the folding and the thioester bond status of the domains are evaluated with a 115 pN pulse. In the case of folding and thioester bond reformation, thioester-intact Cpa domains are detected (~49 nm steps); in the case of having only folding, the full extension of the Cpa domain is observed (~95 nm steps). The buffer used along the experiments contained 50 mM Hepes pH 8.5, 150 mM NaCl, 1 mM EDTA, 10 mM L-ascorbic acid, and was supplemented with 100 mM of methylamine or cystamine for the thioester bond cleavage. To induce thioester bond reformation after cystamine treatment, the same buffer but supplemented with 10 mM of TCEP was added and the force quenched to 4 pN to favor folding and reformation.

**Analysis**

Analysis was done with Igor Pro 8.0 software (Wavemetrics). Recordings were smoothed using a 4th order Savitzky-Golay filter with a box size of 51 points. Step sizes were determined measuring the distance between the peaks of Gaussian fits done on the unfolding steps. Folding probability was calculated as the ratio between the number of unfolded domains and the number of domains able to fold after 100 s at each of the forces tested. Thioester bond cleavage was calculated as the ratio between detected cleavage steps at any of the forces tested, and the number of thioester-intact Cpa domains susceptible to be cleaved. Reformation was calculated as the ratio between thioester-intact Cpa domains detected and the number of cleaved domains registered before. For folding, cleavage and reformation probabilities, a jackknife estimator was used for the calculation of the average probability and the standard deviation.

#### Supplementary Figures

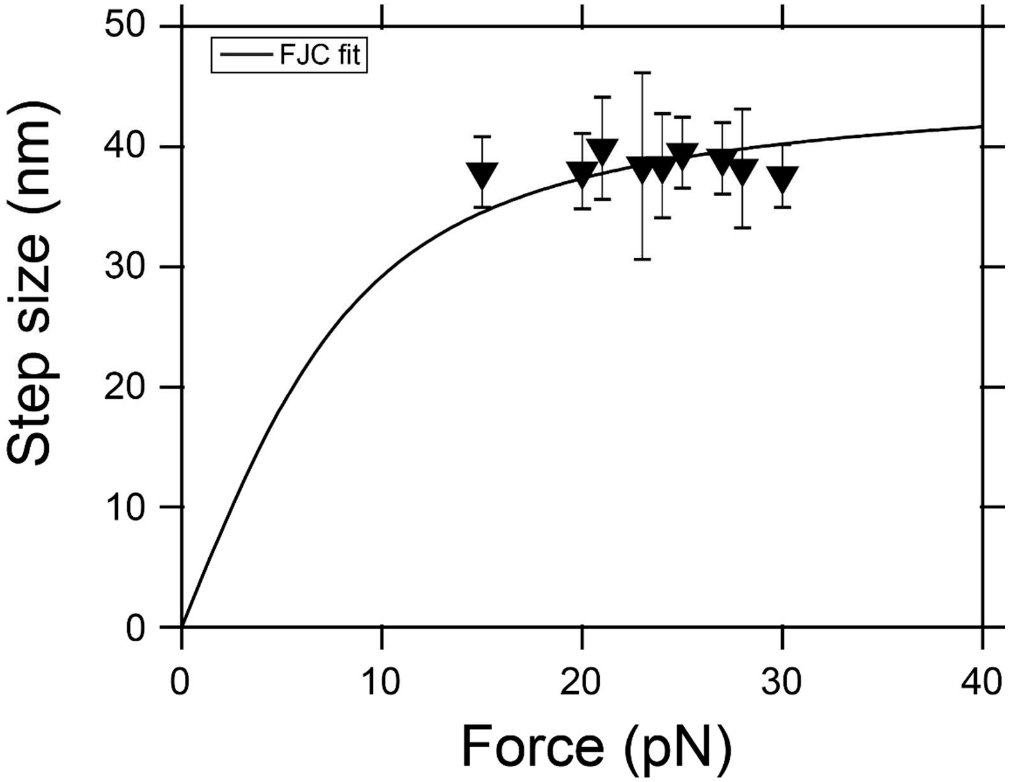


**Supplementary Figure 1. Force-dependent thioester bond cleavage step size.** Cpa thioester bond cleavage step size as a function of force. Data points are the mean±SD of the step size measured at each force (n=29 at 15 pN, n=20 at 20 pN, n=25 at 21 pN, n=34 at 23 pN, n=21 at 24 pN, n=31 at 25 pN, n=18 at 27 pN, n=12 at 28 pN, n=10 at 30 pN). The line is the freely jointed chain fit for polymer elasticity to the Cpa thioester bond-cleavage extensions. The value of contour length $L_c$ was determined subtracting the value for thioester-intact Cpa (49 nm) to thioester-cleaved Cpa (95 nm): $L_c$=95-49=46 nm; Kuhn length value was set to $l_k$=1.1 nm

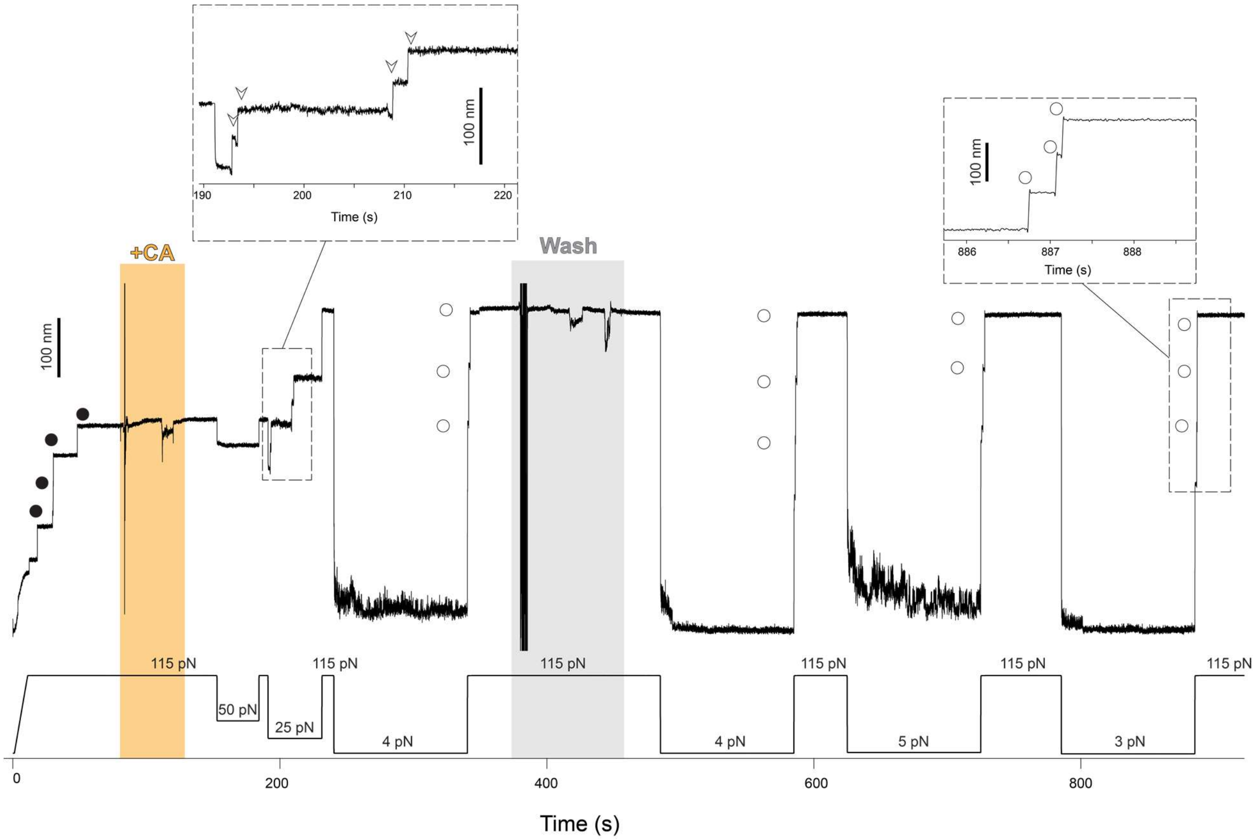


**Supplementary Figure 2. Cystamine permanent blocking of Cpa thioester bond reformation.** Magnetic tweezers force-clamp trajectory of the Cpa polyprotein. After the unfolding of the thioester-intact Cpa domains at 115 pN (circles), the buffer is exchanged and the polyprotein is exposed to a solution containing 100 mM cystamine (+CA). At 115 pN and at 50 pN, no additional extensions are registered as a consequence of thioester bond cleavage, but a drop in force to 25 pN leads to the appearance of four steps which account for the release of the polypeptide sequence trapped by the thioester bonds (empty arrows in the inset). Then, the force is increased again to 115 pN, revealing the complete extension of the molecule. After 100 s at 4 pN and in the presence of cystamine, a 115 pN pulse reveals three ~95 nm steps (empty circles) which correspond with the full extension of Cpa. Cystamine is then removed from the solution, and several consecutive 100 s force quenches (at 4, 5, and 3 pN) followed by 115 pN pulses are applied. These cycles reveal that, after cystamine treatment, Cpa is able to fold but not to reform its thioester bond, as it can be observed from the ~95 nm steps observed (empty circles). After the first 300 s of the experiment, one of the Cpa domains stops folding back as a consequence of oxidative damage[5]. The disturbances observed in the extension during +CA addition (orange block) and washing (gray block) are originated from the movement of buffer volumes in the liquid cell used in the experiments, which transiently alter the measurement.

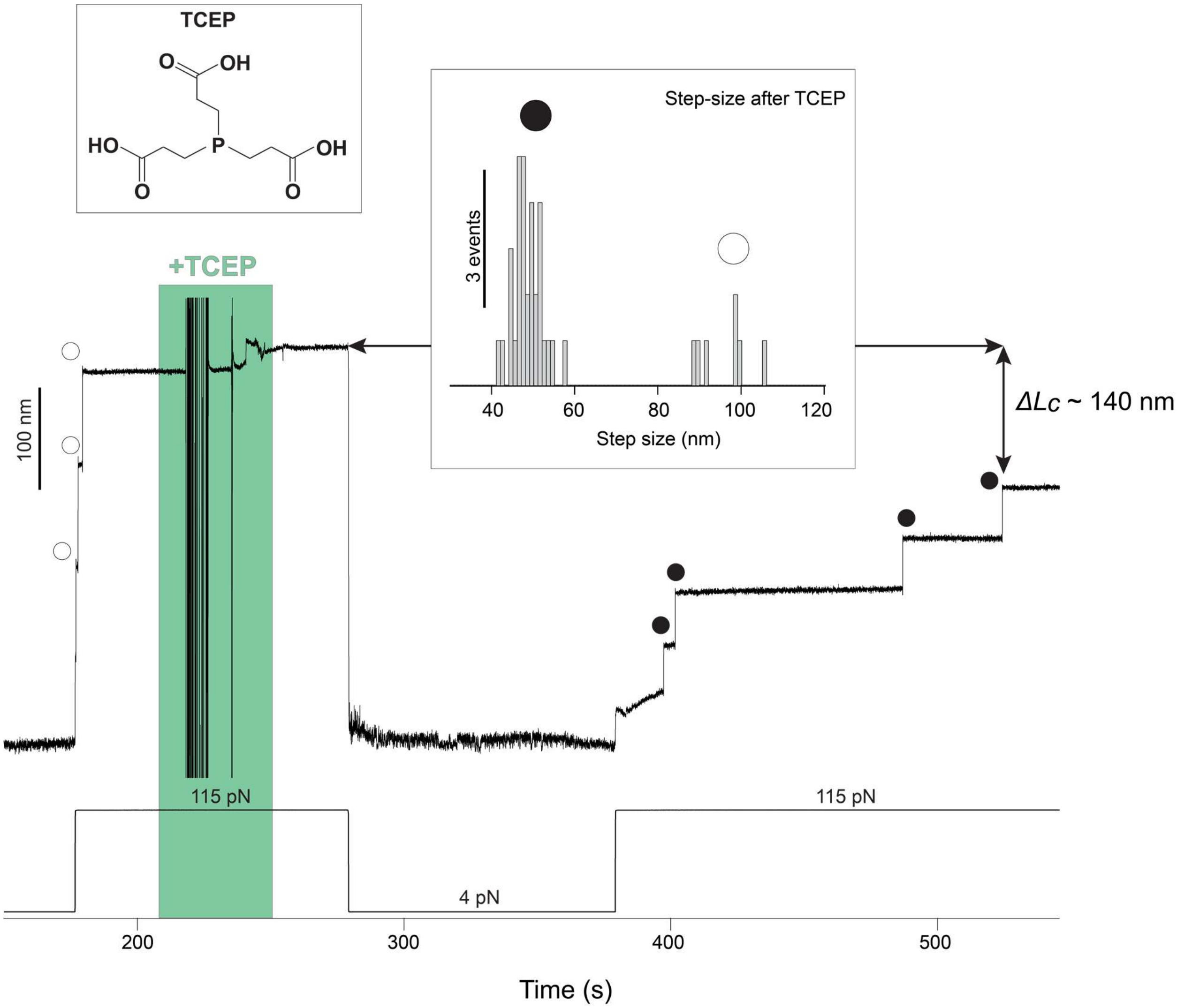


**Supplementary Figure 3. TCEP rescues Cpa thioester bond reformation.** A Cpa polyprotein previously treated with cystamine shows three ~95 nm steps at 115 pN corresponding with the full extension of each of the domains (empty circles). The addition of 10 mM TCEP and 100 s at 4 pN is enough to trigger thioester bond reformation, as it can be observed in the ~49 nm thioester-intact Cpa steps (circles) registered at 115 pN. The fourth domain not observed at the beginning was probably unfolded and its thioester bond intact, since the difference in the final extension between the first 115 pN pulse and the last is ~140 nm, which matches with the expected final extension decrease from three reformation events. Inset histogram shows the two populations of steps observed after TCEP treatment, thioester-intact Cpa (circles, 48.3 ± 3.5 nm, mean±SD, n=32) and thioester-cleaved Cpa (empty circles, 95.7 ± 6.4 nm mean±SD, n=7). The latter full length steps of Cpa after TCEP treatment could be due to cleavage events induced by remaining cystemine which was not completely washed from the experimental liquid cell. The disturbances observed in the extension during +TCEP addition (green block) are originated from the movement of buffer volumes in the liquid cell used in the experiments, which transiently alter the measurement.

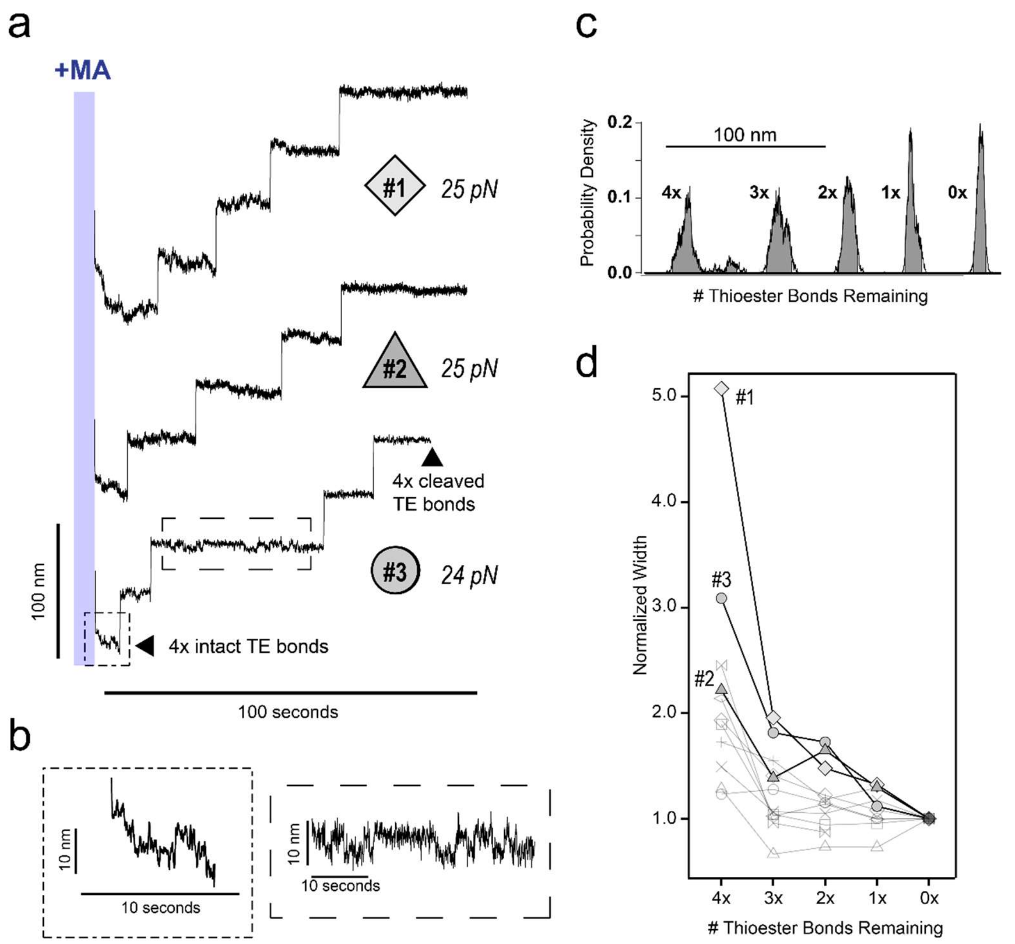


**Supplementary Figure 4. Thioester cleavage follows short lengthwise fluctuations. a)** Representative recordings of Cpa polyprotein after the addition of methylamine and relaxation from 115 pN to 24 and 25 pN. Four thioester bonds are intact at the beginning of each recording and four subsequent thioester cleavage steps follow. **b)** Expansions of the initial collapse trajectory and the equilibrium fluctuations from trajectory #3. **c)** The normalized probability density of each step level in trajectory #1 is shown with the 95% probability width shaded in grey and **d)** plotted as a function of the number of intact thioester bonds remaining for each of 11 single molecule recordings.
